# Loss of sphingomyelin synthase-1 does not cause egg retention or locomotion defects in *Caenorhabditis elegans*

**DOI:** 10.64898/2026.02.10.705178

**Authors:** Wenyue Wang, Xiaoyan Gao, Roger Pocock

**Affiliations:** Development and Stem Cells Program, Monash Biomedicine Discovery Institute and Department of Anatomy and Developmental Biology, Monash University, Melbourne, Victoria 3800, Australia

## Abstract

Sphingomyelin is a critical sphingolipid found in plasma membranes of metazoa that provides structural and communicative functions. Sphingomyelin synthases are key enzymes that generate sphingomyelin but their precise functions in animal development and function are not fully understood. The *Caenorhabditis elegans* model encodes five sphingomyelin synthases (*sms-1-5*). Previously, egg-laying and locomotion phenotypes were observed in an *sms-1(ok2399)* deletion mutant. In this study, we attempted to replicate these findings to enable mechanistic dissection of sphingomyelin function. We indeed found that the *sms-1(ok2399)* mutant exhibited egg-laying and locomotion defects, however, we were unable to rescue this phenotype. Further, we generated two additional *sms-1* deletion mutants (*rp398* and *rp399*) and found that their egg-laying and locomotion behavior is not different to wild-type animals. We suggest that the *sms-1(ok2399)* contains a background mutation that causes behavioral deficits, and that SMS-1 loss does not overtly affect *C. elegans* egg-laying or locomotion.

## INTRODUCTION

Sphingolipids are amphipathic molecules with reported functions in controlling cell adhesion and migration, cell death and cell proliferation (Maceyka and Spiegel, 2014). Given these fundamental functions, studies of sphingolipids and their metabolic pathways have extended to physiological and pathological conditions, such as motor dysfunction, insulin resistance, cancer, cardiovascular disorders, and neurological diseases (Hannun and Obeid, 2018; McCluskey et al., 2022; Pan et al., 2023). Sphingomyelin (SM) is a major structural sphingolipid in cellular membranes, composed of a phosphocholine head and a ceramide backbone (Slotte, 2013). SM interacts with cholesterol to form membrane microdomains (lipid rafts), which act as signalling platforms for cellular communication (Chakraborty and Jiang, 2013). In addition, SM metabolism generates bioactive molecules, such as ceramide, that mediate diverse cellular signalling and regulatory processes (Stith et al., 2019).

SM levels are tightly controlled by a network of enzymes, including sphingomyelin synthases and sphingomyelinases, with ceramide acting both as a SM precursor and breakdown product (Kuo and Hla, 2024). Disruption of SM metabolism, often accompanied by altered ceramide levels, has been implicated in motor neuron diseases, where it may contribute to neuronal degeneration by affecting membrane organization, lipid raft–associated signaling, and cellular stress responses (McCluskey et al., 2022). Despite these associations, it remains unclear how individual SM metabolic enzymes specifically influence motor neuron function *in vivo*.

We previous showed that the nematode *Caenorhabditis elegans* is an excellent genetically tractable model to investigate sphingolipid functions (Wang et al., 2023). The *C. elegans* genome encodes five sphingomyelin synthases (Guzman et al., 2025). A previous study reported that SMS-1 loss impairs motor neuron–dependent behaviors, such as locomotion and egg-laying, but the underlying mechanisms remained unresolved (Hao et al., 2017). In this study, we re-examined the potential role of SMS-1 in *C. elegans* egg-laying and locomotion by generating independent *sms-1* deletion alleles using CRISPR/Cas9, and performing quantitative behavioral assays and transgenic rescue. Our results reveal that the egg retention and locomotion phenotypes previously observed in the *sms-1(ok2399)* deletion mutant are not caused by SMS-1 loss. Our findings underscore the importance of rigorous genetic validation when analysing gene knockouts.

## RESULTS

### The *sms-1(ok2399)* deletion allele causes egg retention and reduced locomotion

A previous study identified egg-laying and locomotion phenotypes in *sms-1(ok2399)* animals, without dissecting the underlying mechanism (Hao et al., 2017). To investigate this, we attempted to replicate these behavioral phenotypes after backcrossing the *sms-1(ok2399)* allele with wild-type males five times (Fig. 1).

**Fig. 1.**
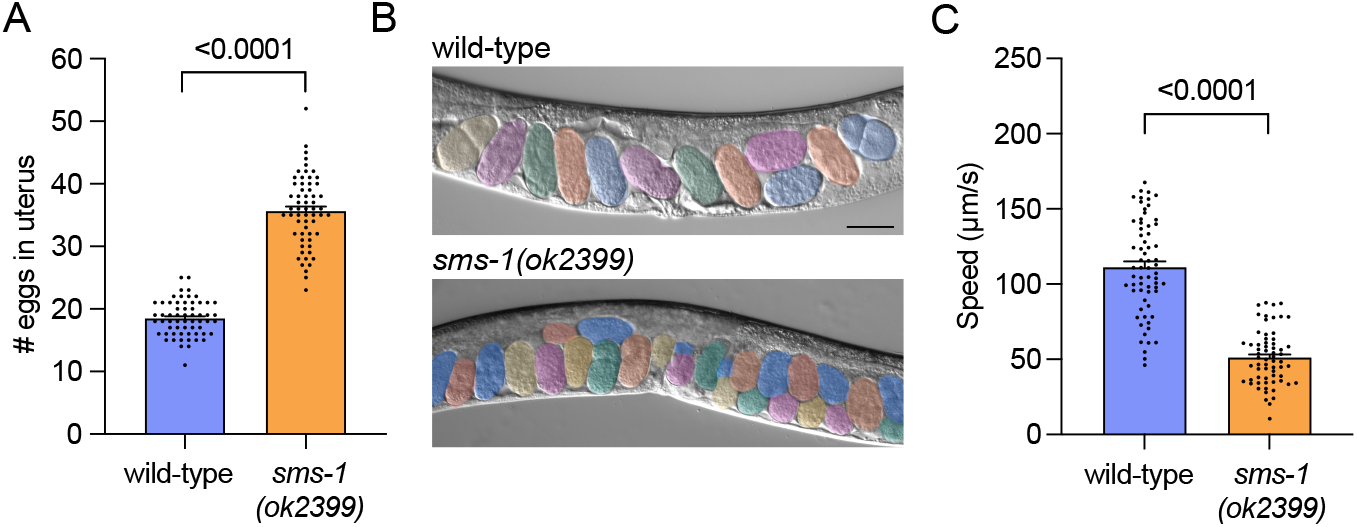
*sms-1(ok2399)* animals exhibit excess egg retention and reduced locomotion. (A-B) Quantification (A) and DIC micrographs (B) of wild-type and *sms-1(ok2399)* egg retention in one-day old hermaphrodites. Eggs are pseudocolored. Worm images are oriented anterior to the left, ventral down. n = 60. Scale bar, 50 µm. (C) Quantification of wild-type and *sms-1(ok2399)* locomotory speed (distance travelled/time) in one-day old hermaphrodites. n = 64-65. Statistical significance was assessed using an unpaired t test (A and C). Error bars indicate SEM. Significance was considered at p<0.05.

In *C. elegans*, egg-laying is a highly regulated, rhythmic behavior that is controlled by multiple neuronal and environmental inputs (Fenk and de Bono, 2015; Zhang et al., 2008). Young adult hermaphrodites typically retain ∼15 fertilized eggs in their uterus. When regulatory inputs are disrupted, the egg-laying rate may increase or decrease, thus altering the number of eggs retained in the uterus at a defined developmental timepoint (Scharf et al., 2021). We cultivated wild-type and *sms-1(ok2399)* animals in parallel, and counted the number of eggs retained in the uterus of one-day old adult hermaphrodites (36 hours after the mid-L4 larval stage of development). Consistent with previous data, we found that *sms-1(ok2399)* one-day old adult hermaphrodites retained more eggs (∼35) than wild-type animals (∼18) (Fig. 1A-B). To confirm that SMS-1 also controls motor function, we measured locomotion speed of wild-type and *sms-1(ok2399)* one-day adult hermaphrodites (24 hours after the mid-L4 larval stage of development). We found that *sms-1(ok2399)* mutants exhibit reduced locomotion (∼51µm/s) compared with wild-type animals (∼111 µm/s), confirming previous findings (Fig. 1C). Together, these results show that *sms-1(ok2399)* animals exhibit defects in egg-laying and locomotion, consistent with the previously report.

### Overexpressing *sms-1* fails to rescue egg retention and locomotion defects

The *sms-1* gene is predicted to generate a protein containing 6 transmembrane domains (Hao et al., 2017). The *sms-1(ok2399)* deletion is likely a loss of function allele as it is predicted to disrupt splicing and cause a frameshift, resulting in loss of the protein sequence from transmembrane domain 3 onward (Hao et al., 2017).

To assess whether loss of *sms-1* underlies the egg retention and locomotion defects, we attempted phenotypic rescue by reintroducing *sms-1* genomic DNA driven by its endogenous promoter (2 kb upstream of the start codon) into *sms-1(ok2399)* mutant animals (Fig. 2A-C). However, analysis of three independent transgenic lines showed that *sms-1* overexpression failed to rescue either the egg retention or locomotion defects (Fig. 2B-C).

**Fig. 2.**
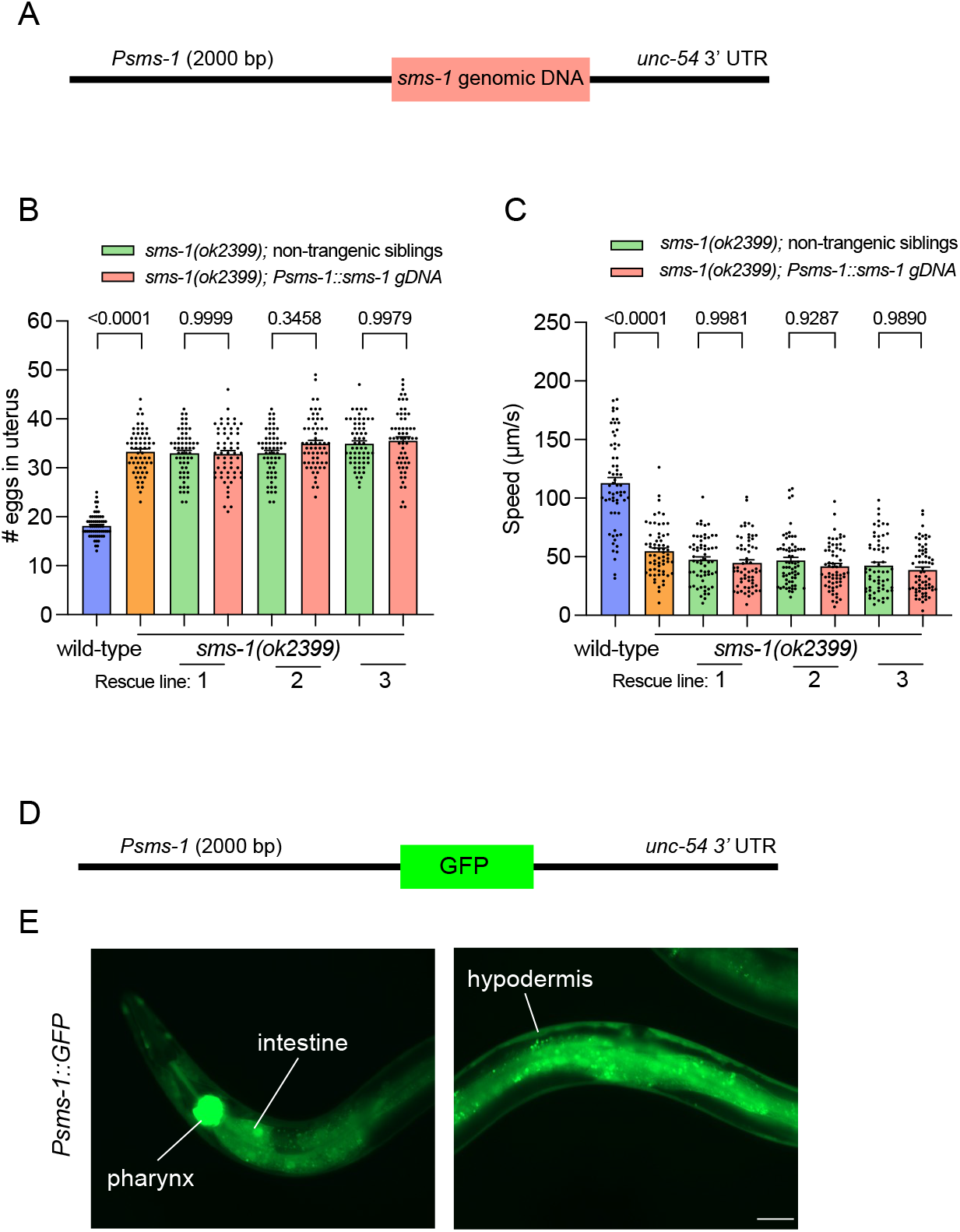
Transgenic *sms-1* expression fails to rescue the *sms-1(ok2399)* egg retention and locomotion phenotypes. (A) Schematic of the *Psms-1::sms-1*-genomic DNA rescue construct used to rescue the egg retention and locomotion phenotypes of *sms-1(ok2399)* animals. *Psms-1* = 2000bp *sms-1* promoter region; *sms-1* genomic DNA = 4679 bp genomic DNA; *unc-54* 3’ UTR = untranslated region. (B) Quantification of wild-type, *sms-1(ok2399)*, and *sms-1(ok2399); Psms-1::sms-1 gDNA* transgenic animal egg retention in one-day old hermaphrodites. Green bars, non-transgenic siblings; pink bars, transgenic animals. n = 60. (C) Quantification of wild-type, *sms-1(ok2399)*, and *sms-1(ok2399); Psms-1::sms-1 gDNA* transgenic animal locomotory speed (distance travelled/time) in one-day old hermaphrodites. Green bars, non-transgenic siblings; pink bars, transgenic animals. n = 61-65. (D) Schematic of the *Psms-1::GFP* fluorescent reporter construct used to examine the where the *sms-1* promoter used in the rescue experiments (A-C) drives expression. (E) Expression of the *Psms-1::GFP* fluorescent reporter in wild-type animals. GFP expression is detected in the pharynx and intestine (left panel) and hypodermis (right panel). Scale bar, 50 µm. Statistical significance was assessed using one-way ANOVA followed by a Tukey’s multiple comparisons test (B and C). Error bars indicate SEM. Significance was considered at p<0.05.

RNA sequencing analysis shows that *sms-1* is predominantly expressed in the hypodermis, with lower-level of expression all other tissues (Cao et al., 2017; Packer et al., 2019). To confirm that the endogenous *sms-1* promoter we used for the rescue experiments drives expression in the correct tissues, we generated a GFP reporter driven by the same promoter (Fig. 2D-E). This reporter recapitulated the expected *sms-1* expression pattern, with strong expression in the hypodermis and other tissues, including the intestine and pharynx (Fig. 2E). The lack of rescue suggested to us that the egg retention and locomotion defects observed in *sms-1(ok2399)* may not be caused by *sms-1* loss and potentially by a background mutation.

### CRISPR/Cas9-generated *sms-1* deletion alleles (*rp398* and *rp399*) exhibit wild-type egg-laying and locomotion

To independently assess the function of SMS-1 in egg-laying and locomotion, we generated two independent *sms-1* alleles using CRISPR/Cas9 (Fig. 3A). In contrast to the *ok2399* allele, which deletes 620 bp spanning exons 3–4, the newly generated alleles, *rp398* and *rp399*, contain larger deletions of 3163 bp and 3086 bp, respectively, that remove the majority of the *sms-1* coding region (Fig. 3A). These deletions are thus predicted null alleles. We next assessed egg-laying and locomotion behaviors in all three *sms-1* deletion mutants in parallel with wild-type animals. We found that the *sms-1(rp398)* and *sms-1(rp399)* animals exhibited wild-type egg retention and locomotion defects, while *sms-1(ok2399)* were defective. Together, these results indicate that the phenotypes previously associated with the *sms-1(ok2399)* allele are not due to loss of *sms-1* function. Thus, we suggest that future studies of SMS-1 function should utilise the *rp398* and *rp399* alleles we generated.

**Fig. 3.**
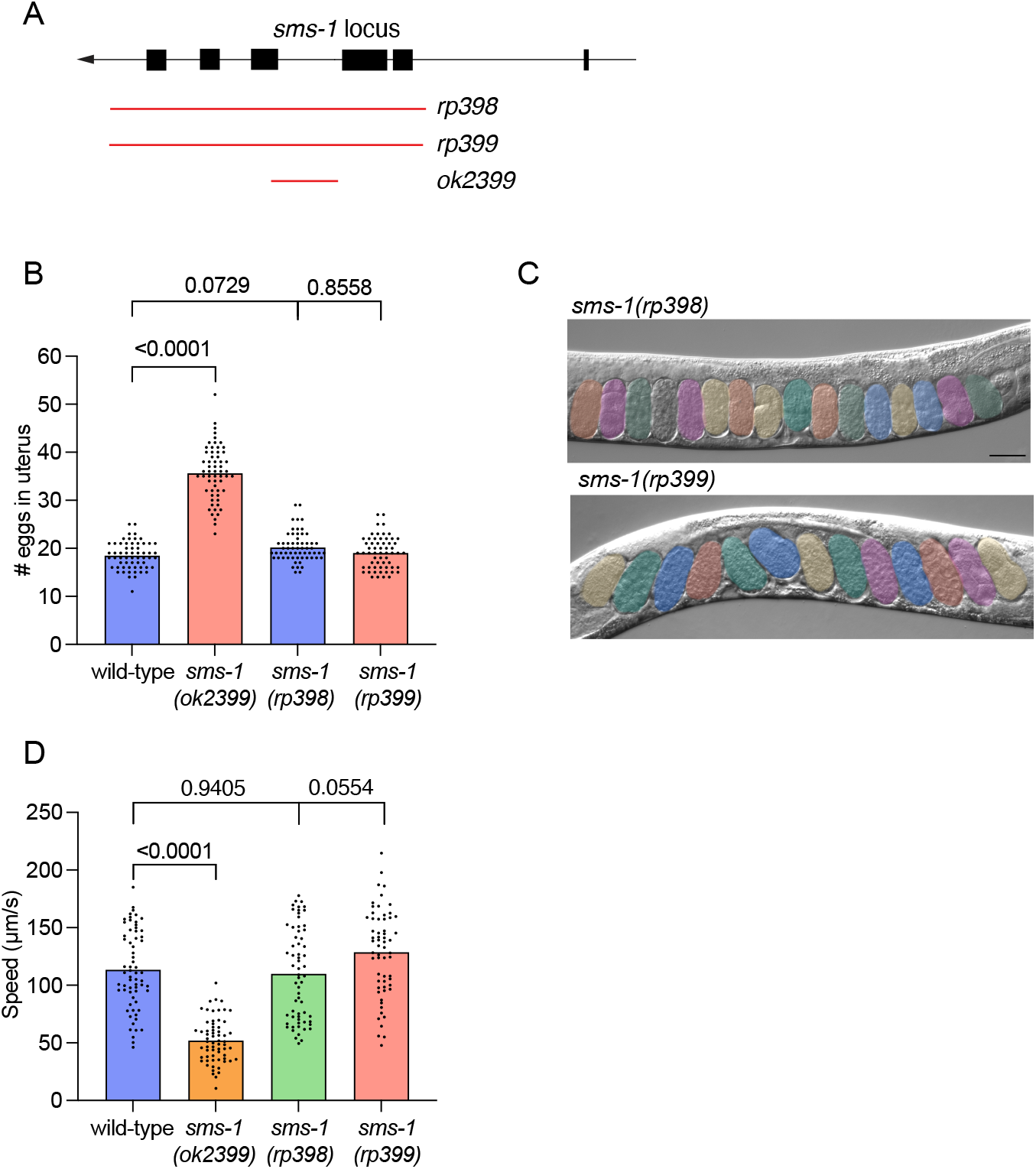
CRISPR/Cas9-generated *sms-1* deletion alleles (*rp398* and *rp399*) do not phenocopy the *ok2399* egg retention and locomotion phenotypes. (A) Structure of the *sms-1* locus. *ok2399* = 620 bp deletion; *rp398* = 3163 bp deletion; *rp399* =3086 bp deletion. Black boxes, coding regions; black lines, introns; red lines, deletion alleles. (B) Quantification of wild-type and *sms-1(ok2399, rp398* and *rp399)* egg retention in one-day old hermaphrodites. (C) DIC micrographs of *sms-1(rp398* and *rp399)* egg retention in one-day old hermaphrodites (compare with wild-type and *sms-1(ok2399)* images in Fig 1B). Eggs are pseudocolored. Worm images are oriented anterior to the left, ventral down. n = 60. Scale bar, 50 µm. (D) Quantification of wild-type and *sms-1(ok2399, rp398* and *rp399*) locomotory speed (distance travelled/time) in one-day old hermaphrodites. n = 61-64. Statistical significance was assessed using one-way ANOVA followed by a Tukey’s multiple comparisons test (B and D). Error bars indicate SEM. Significance was considered at p<0.05.

## DISCUSSION

Our results provide a rigorous reassessment of the role of *sms-1* in regulating egg-laying and locomotion in *C. elegans*. Consistent with a previous report, the *sms-1(ok2399)* allele exhibited excess egg retention and impaired locomotion. However, overexpression of the *sms-1* gene driven by its endogenous promoter did not restore normal egg-laying or locomotion, despite confirming promoter activity in the pharynx, intestine, and hypodermis. To further test whether *sms-1* loss underlies these behavioral phenotypes, we generated two independent CRISPR/Cas9 deletion alleles, *rp398* and *rp399*, which remove the majority of the coding sequence and are predicted nulls. These alleles exhibited normal egg-laying and locomotion, in contrast to the *ok2399* allele. Our findings indicate that the phenotypes associated with the *ok2399* allele are not a consequence of *sms-1* loss, indicating that *ok2399* may harbor additional background mutations or allele-specific effects that contribute to the observed defects.

Our study underscores the importance of using multiple independent alleles and genetic validation when linking specific genes to phenotypes. We suggest that future studies investigating SMS-1 function in controlling SM biology, including lipidomics, should employ our CRISPR/Cas9-generated *sms-1* alleles which we will deposit at the *Caenorhabditis Genetics Center*.

## MATERIALS AND METHODS

### *C. elegans* strains and culture

*C. elegans* hermaphrodites were maintained according to standard protocols at 20°C on nematode growth medium (NGM) plates with *Escherichia coli* (OP50) bacteria as a food source. The wild-type strain used was Bristol, N2. Mutant strains were backcrossed to N2 five times and animals were well-fed for at least two generations before performing experiments. A list of the strains and plasmids used in this study, and source data is provided in Table S1.

### CRISPR-Cas9 genome editing

To generate *sms-1* deletion mutants, adult wild-type hermaphrodites were microinjected with Cas9 protein, tracrRNA, and two crRNAs: 5’ crRNA (CCCTTTAACAGTTGACTCAT), 3’ crRNA (GATGATGTTCCCGGGTGCTT) from IDT. *sms-1* deletions were identified by PCR and confirmed by Sanger sequencing. The *sms-1(rp398)* and *sms-1(rp399)* alleles generated are 3163 bp and 3086 bp deletions, respectively. Both deletion alleles remove most of the *sms-1* gene (Fig. 3A). Each allele was backcrossed to wild-type males prior to analysis.

### *C. elegans* transgenic strain generation

Reporter gene constructs were generated by PCR amplifying DNA elements and cloning them into Fire vectors. The constructs were injected into young adult hermaphrodites using standard methods.

#### sms-1p::gfp reporter construct

A 2,000 bp sequence upstream of the *sms-1* start codon was amplified from *C. elegans* genomic DNA using forward (5′ TGATTACGCCAAGCTTAGAGGGCTCTGGAGAAAAACC -3′) and reverse (5′-CCAATCCCGGGGATCCGTTATGTCACACGGTGTTCG -3′) oligonucleotides incorporating *HindIII-BamHI* restriction sites. The *HindIII-BamHI-*digested promoter fragment was ligated into *HindIII-BamHI-*digested pPD95.75 vector, resulting in *Psms-1::gfp*, which was injected at 20 ng/μl, with 5 ng/µl *Pmyo-2::mCherry* vector into wild type animals.

#### Psms-1::sms-1 genomic DNA extrachromosomal array rescue construct

*sms-1(isoform a)* gDNA (4679 bp) was amplified from genomic DNA using forward (5′ AGGACCCTTGGCTAGCATGAAAATGTCTTGGAATCATCAA -3′) and reverse (5′-GATATCAATACCATGGTCATTCGAAAGCAGGTCGTG -3′) oligonucleotides incorporating *NheI-NcoI* restriction sites. The *NheI-NcoI-*digested promoter fragment was inserted using the In-Fusion HD Cloning Kit (Takara Bio) into *NheI-NcoI-*digested *ges-1p::asah-1 cDNA* vector (to remove *asah-1* cDNA). A 2,000 bp sequence upstream of the *sms-1* start codon was then amplified from *C. elegans* genomic DNA with forward (5′-TGATTACGCCAAGCTTAGAGGGCTCTGGAGAAAAACC -3′) and reverse (5′-CCAATCCCGGGGATCCGTTATGTCACACGGTGTTCG-3′) oligonucleotides from incorporating *HindIII-BamHI* restriction sites. The *Pges-1::sms-1 gDNA* plasmid was digested using *HindIII-BamHI* to remove the *ges-1* promoter and the *sms-1* promoter was inserted using the In-Fusion HD Cloning Kit (Takara Bio). The resultant *Psms-1::sms-1 genomic DNA* plasmid was injected at 50 ng/μl, with 5 ng/μl *Pmyo-2::mCherry* vector into *sms-1(ok2399)* animals.

### Locomotion analysis

Worm locomotion was analysed at room temperature on one-day adults using WormLab imaging (MBF Bioscience). NGM plates used for tracking were freshly seeded with 20 µl OP50 10 min before use. To standardize acclimation time and food conditions across groups, plates were analysed in a strictly interleaved rotating order until six plates were analysed per group. Six animals were placed on each plate and their tracks were recorded for 1 min.

### Egg retention analysis

The number of eggs in utero were counted in age-synchronized hermaphrodites. Worms were picked at the mid-L4 stage based on the morphology of the vulval invagination. 36h later, individual worms were lysed in 20 µl of a freshly prepared lysis buffer (1:1 mixture of household bleach and NaOH). Once hermaphrodites dissolved, the released eggs were counted under a dissecting microscope.

### Microscopy / Expression analysis

Worms were imaged at the L4 stage to determine the expression pattern of *Psms-1::gfp*. Worms were anaesthetized using levamisole (0.1 ng/μl) and mounted on 5% agarose pads on glass slides. Fluorescence images were acquired using a Zeiss Axio Imager M2 and Zen software tools. Figures were prepared using ImageJ and Adobe Illustrator.

### Statistics and reproducibility

Statistical analyses were conducted in GraphPad Prism (v9.5). Significance (P < 0.05) was determined via unpaired Student’s t-test for two-group comparisons, or ANOVA (with Dunnett’s or Tukey’s post-hoc tests) for multiple conditions. All experiments were performed with at least three independent biological replicates. No data were excluded, and investigators were blinded to genotype during analysis.

## Acknowledgments

We thank members of the Pocock Laboratory for comments on the manuscript. Some strains were provided by the *Caenorhabditis* Genetics Center (University of Minnesota), which is funded by NIH Office of Research Infrastructure Programs (P40 OD010440).

## Author contributions

Conceptualization, W.W. and R.P.; Methodology, W.W., X.G. and R.P.; Investigation, W.W., X.G. and R.P.; Writing – Original Draft, W.W.; Writing – Review & Editing, W.W., X.G. and R.P.; Funding Acquisition, R.P.; Resources, R.P; Supervision, W.W. and R.P.

## Data and materials availability

All data is available in the main text and Source Data in Table S1. There are no accession codes, unique identifiers, or weblinks in our study and no restrictions on data availability. Materials will be available upon request from the Pocock laboratory.

## SUPPLEMENTARY INFORMATION

**Table S1.Reagent information and Source Data**

## REFERENCES

Cao, J., Packer, J. S., Ramani, V., Cusanovich, D. A., Huynh, C., Daza, R., Qiu, X., Lee, C., Furlan, S. N., Steemers, F. J., et al. (2017). Comprehensive single-cell transcriptional profiling of a multicellular organism. Science 357, 661–667.

Chakraborty, M. and Jiang, X. C. (2013). Sphingomyelin and its role in cellular signaling. Adv Exp Med Biol 991, 1–14.

Fenk, L. A. and de Bono, M. (2015). Environmental CO2 inhibits Caenorhabditis elegans egg-laying by modulating olfactory neurons and evokes widespread changes in neural activity. Proc Natl Acad Sci U S A 112, E3525–3534.

Girard, L. R., Fiedler, T. J., Harris, T. W., Carvalho, F., Antoshechkin, I., Han, M., Sternberg, P. W., Stein, L. D. and Chalfie, M. (2007). WormBook: the online review of Caenorhabditis elegans biology. Nucleic Acids Res 35, D472–475.

Guzman, G., Jahn, H., Farley, S. E., Kyle, J. E., Bramer, L. M., Hoetzl, S., van den Dikkenberg, J., Hermansson, M., Holthuis, J. C. M. and Tafesse, F. G. (2025). Characterization of Caenorhabditis elegans sphingomyelin synthases through heterologous expression. J Biol Chem 301, 110300.

Hannun, Y. A. and Obeid, L. M. (2018). Author Correction: Sphingolipids and their metabolism in physiology and disease. Nat Rev Mol Cell Biol 19, 673.

Hao, L., Ben-David, O., Babb, S. M., Futerman, A. H., Cohen, B. M. and Buttner, E. A. (2017). Clozapine Modulates Glucosylceramide, Clears Aggregated Proteins, and Enhances ATG8/LC3 in Caenorhabditis elegans. Neuropsychopharmacology 42, 951–962.

Kuo, A. and Hla, T. (2024). Regulation of cellular and systemic sphingolipid homeostasis. Nat Rev Mol Cell Biol 25, 802–821.

Maceyka, M. and Spiegel, S. (2014). Sphingolipid metabolites in inflammatory disease. Nature 510, 58–67.

McCluskey, G., Donaghy, C., Morrison, K. E., McConville, J., Duddy, W. and Duguez, S. (2022). The Role of Sphingomyelin and Ceramide in Motor Neuron Diseases. J Pers Med 12.

Packer, J. S., Zhu, Q., Huynh, C., Sivaramakrishnan, P., Preston, E., Dueck, H., Stefanik, D., Tan, K., Trapnell, C., Kim, J., et al. (2019). A lineage-resolved molecular atlas of C. elegans embryogenesis at single-cell resolution. Science 365.

Pan, X., Dutta, D., Lu, S. and Bellen, H. J. (2023). Sphingolipids in neurodegenerative diseases. Front Neurosci 17, 1137893.

Scharf, A., Pohl, F., Egan, B. M., Kocsisova, Z. and Kornfeld, K. (2021). Reproductive Aging in Caenorhabditis elegans: From Molecules to Ecology. Front Cell Dev Biol 9, 718522.

Slotte, J. P. (2013). Biological functions of sphingomyelins. Prog Lipid Res 52, 424–437.

Stith, J. L., Velazquez, F. N. and Obeid, L. M. (2019). Advances in determining signaling mechanisms of ceramide and role in disease. J Lipid Res 60, 913–918.

Wang, W., Sherry, T., Cheng, X., Fan, Q., Cornell, R., Liu, J., Xiao, Z. and Pocock, R. (2023). An intestinal sphingolipid confers intergenerational neuroprotection. Nat Cell Biol 25, 1196–1207.

Zhang, M., Chung, S. H., Fang-Yen, C., Craig, C., Kerr, R. A., Suzuki, H., Samuel, A. D., Mazur, E. and Schafer, W. R. (2008). A self-regulating feed-forward circuit controlling C. elegans egg-laying behavior. Curr Biol 18, 1445–1455.

